# An In Vitro Medium for Modeling Gut Dysbiosis Associated with Cystic Fibrosis

**DOI:** 10.1101/2023.08.01.551570

**Authors:** Kaitlyn E. Barrack, Thomas H. Hampton, Rebecca A. Valls, Sarvesh V. Surve, Timothy B. Gardner, Julie L. Sanville, Juliette C. Madan, George A. O’Toole

## Abstract

The gut physiology of pediatric and adult persons with cystic fibrosis (pwCF) is altered relative to healthy persons. The CF gut is characterized, in part, as having excess mucus, increased fat content, acidic pH, increased inflammation, increased antibiotic perturbation and the potential for increased oxygen availability. These physiological differences shift nutritional availability and the local environment for intestinal microbes, thus likely driving significant changes in microbial metabolism, colonization and competition with other microbes. The impact of any specific change in this physiological landscape is difficult to parse using human or animal studies. Thus, we have developed a novel culture medium representative of the CF gut environment, inclusive of all the aforementioned features. This medium, called CF-MiPro, maintains CF gut microbiome communities, while significantly shifting non-CF gut microbiome communities toward a CF-like microbial profile, characterized by low Bacteroidetes and high Proteobacteria abundance. This medium is able to maintain this culture composition for up to 5 days of passage. Additionally, microbial communities passaged in CF-MiPro produce significantly less immunomodulatory short chain fatty acids (SCFA), including propionate and butyrate, than communities passaged in MiPro, a culture medium representative of healthy gut physiology, confirming not only a shift in microbial composition but altered community function. Our results support the potential for this in vitro culture medium as a new tool for the study of gut dysbiosis in CF.

**Importance:** Cystic fibrosis is an autosomal recessive disease that disrupts ion transport at mucosal surfaces, leading to mucus accumulation and altered physiology of both the lungs and the intestines, among other organs, with the resulting altered environment contributing to an imbalance of microbial communities. Culture media representative of the CF airway have been developed and validated; however, no such medium exists for modeling the CF intestine. Here, we develop and validate a first-generation culture medium inclusive of features that are altered in the CF colon. Our findings suggest this novel medium, called CF-MiPro, as a maintenance medium for CF gut microbiome samples and a flexible tool for studying key drivers of CF-associated gut dysbiosis.

## Introduction

Cystic fibrosis (CF) is a hereditary condition caused by mutations in the gene encoding the cystic fibrosis transmembrane conductance regulator (CFTR) protein. Loss of CFTR function leads to disrupted chloride and bicarbonate secretion and an imbalance in hydration of mucosal surfaces (1). Clinical consequences of CF include, but are not limited to, accumulation of thick mucus at epithelial surfaces, impaired mucociliary clearance due to changes in fluid and ion fluxes, exocrine dysfunction, obstruction of the airway and intestine, inflammation, and increased susceptibility to infection (2) resulting in significant multisystem morbidity and premature mortality.

While much attention has been given to the effects of CF on the airways, there is substantial evidence indicating that the intestine is also affected by this disease, as CFTR activity is essential for proper gut function (3, 4). The changes in CF intestinal physiology result in malnutrition, malabsorption of fats and vitamins, increased acidity, and poor linear growth (3, 5, 6). The dehydrated lumen of the CF gut accumulates mucus, which can significantly slow gastrointestinal motility (7), increasing the risk of delayed meconium passage and meconium ileus in newborns (3, 5). Increased mucus has also been associated with overgrowth of select organisms (1, 8) which can lead to severe gastrointestinal (GI) symptoms and weight loss. Further, the CF gut is characterized by an increase in inflammation, quantified through inflammatory measures, including calprotectin (1, 3, 5, 9–13) and intestinal morphological abnormalities (3, 10, 13). Inflammation further exacerbates the release of reactive oxygen species (ROS) and hemoglobin-carrying oxygen, resulting in oxidative stress and elevated oxygen levels, which in turn contribute to ongoing inflammation within the intestine (14) with consequences for systemic inflammation.

The altered CF intestinal milieu fosters a distinct microbiota. Starting in early life, children with CF (cwCF) have significantly reduced alpha diversity based on stool 16S rRNA amplicon library (5, 14, 15) and metagenomic sequencing (5, 16). The relative age of the fecal microbiota is delayed in cwCF compared to healthy controls (5), and microbes associated with immune training are decreased in CF over the first year of life (15). Considering that the development of the gut microbiota occurs over the first three years of life, and the interaction of the microbiota with the host is essential for immune training within this timeframe, dysbiosis in this critical window has the potential for increased susceptibility to infection, inflammation, allergy, and metabolic disorders (17). Published studies indicate that this dysbiosis persists into later childhood, adolescence, and adulthood for individuals with CF (5, 14, 15, 18, 19). Consequently, this enduring dysbiosis can have adverse effects on persons with CF (pwCF) throughout their lifetime. Notable taxonomic differences observed in cwCF include a decrease in certain immune-modulating bacterial genera of the phyla Bacteroidetes (*Bacteroides*), Firmicutes (*Faecalibacterium*), Actinobacteria (*Bifidobacterium*) and Verrucomicrobia (*Akkermansia*), all positively associated with gut health (5, 14, 15, 20–22). There is a consistent increase in Proteobacteria (*Escherichia coli/Shigella* spp.) and select Firmicutes (*Veillonella, Clostridium*) for pwCF (5, 11, 20–22). These signatures of microbial dysbiosis are also observed in other intestinal conditions, such as inflammatory bowel diseases (23, 24) and in individuals chronically exposed to inorganic arsenic (25–27). Thus, investigation of CF-associated microbial dysbiosis likely has application to other diseases with a similar inflammatory intestinal environment.

A published study has shown that a CFTR mutation is sufficient to alter the GI microbiome in a germ-free CF mouse model (28), suggesting that CF gut physiology is causative of microbial dysbiosis, rather than correlative. A few additional studies have investigated gut microbial dysbiosis in CF animal models (28–32), however it is difficult to identify the role for CF-relevant physiological features, individually or in combination, on microbial dysbiosis in an animal model due to the significant involvement of the host immune system and the likelihood that multiple factors are impacting the microbial communities. Identifying which CF-relevant physiological feature(s) drive the observed dysbiosis associated with this disease is a critical first step in developing strategies to restore intestinal homeostasis.

In this study, we sought to develop and validate a first-generation culture medium that resembles key features of the CF colon. Development and validation of a culture medium representative of the CF colon will allow for in vitro experimentation under more physiologically-relevant conditions. Based on a thorough literature review and consultation with physicians, we modified an existing gut microbiome medium (33) to reflect CF-relevant features. We aimed to generate a first-generation CF colon medium which incorporates sources of bile, mucin, fat, oxidative stress, antibiotics and alternative nutrient sources reported to be increased in states of intestinal inflammation (e.g., nitrate, sulfate, formate) (13, 34–36). We call this new medium CF-MiPro. Using this medium, we show that CF gut microbiota structure is largely maintained at the phylum level and short chain fatty acid (SCFA) profiles are unaffected. Conversely, we can use CF-MiPro to drive CF-relevant shifts in nonCF clinical samples in terms of both composition and function. This novel medium offers a flexible and inexpensive platform to study CF gut dysbiosis in the laboratory.

## Results

### Formulation of CF-MiPro

To develop an in vitro medium representative of the nutritional conditions of the CF colon, we first reviewed the literature to determine differential physiological features described for the CF GI tract, typically via the analysis of stool, and their relevant concentration ranges. Based on these observed ranges, we categorized a “low” and “median” concentration for each feature. In early experiments including all features at the “high” concentration resulted in significant reduction in the population of many microbes (data not shown).

We modified the concentrations of these features in the recipe for MiPro, an in vitro medium designed to mimic the conditions of the healthy colon (33). We call these CF-like media “low-CF-MiPro” and “median-CF-MiPro”. The features and respective concentrations in each medium (low, median) are as follows (see also **Table 1**): sulfate (0.5, 1 mM), a precursor of H2S, which is increased in cases of gut inflammation (34, 35); nitrate (0.5, 1 mM), a by-product of the host inflammatory response (37); formate (0.5, 1 mM), a microbially-derived product increased in the inflamed gut (36); glycerol (0.5, 1%), a marker of increased fat (38, 39); Bactrim (1, 10 uM), an antibiotic commonly prescribed to cwCF in our local cohort at Dartmouth and more broadly (40); H2O2 (1, 10 uM), a measure of inflammation-derived oxidative stress (41, 42); pH (6–7), to represent the general range of acidity in the CF and healthy colons (43, 44); mucin (6, 8 g/L), to represent the thicker mucus layer secondary to CFTR dysfunction (3, 45); and bile salts (1, 2 g/L), to represent the impaired uptake of bile salts in CF (46, 47). We use low-CF-MiPro and median-CF-MiPro in the experiments outlined below and use MiPro as a control in all studies.

**Table 1.**
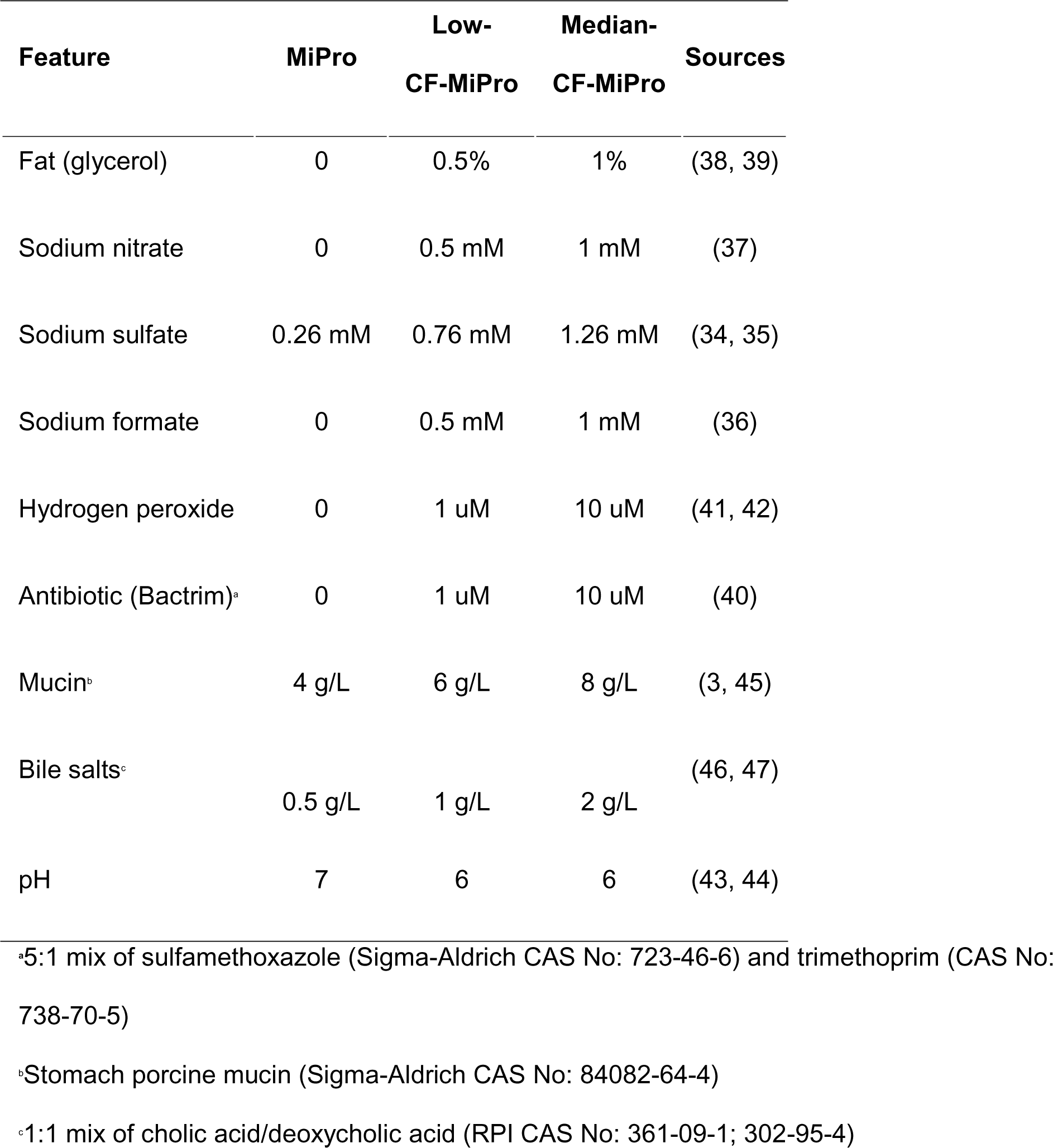
Features and respective concentrations used in MiPro, low-CF-MiPro and median-CF-MiPro.

| Feature | MiPro | Low-CF-MiPro | Median-CF-MiPro | Sources |
| --- | --- | --- | --- | --- |
| Fat (glycerol) | 0 | 0.5% | 1% | (38, 39) |
| Sodium nitrate | 0 | 0.5 mM | 1 mM | (37) |
| Sodium sulfate | 0.26 mM | 0.76 mM | 1.26 mM | (34, 35) |
| Sodium formate | 0 | 0.5 mM | 1 mM | (36) |
| Hydrogen peroxide | 0 | 1 uM | 10 uM | (41, 42) |
| Antibiotic (Bactrim)<sup>a</sup> | 0 | 1 uM | 10 uM | (40) |
| Mucin<sup>b</sup> | 4 g/L | 6 g/L | 8 g/L | (3, 45) |
| Bile salts<sup>c</sup> | 0.5 g/L | 1 g/L | 2 g/L | (46, 47) |
| pH | 7 | 6 | 6 | (43, 44) |

<sup>a</sup>5:1 mix of sulfamethoxazole (Sigma-Aldrich CAS No: 723-46-6) and trimethoprim (CAS No: 738-70-5)
<sup>b</sup>Stomach porcine mucin (Sigma-Aldrich CAS No: 84082-64-4)
<sup>c</sup>1:1 mix of cholic acid/deoxycholic acid (RPI CAS No: 361-09-1; 302-95-4)

### Analysis of inoculum used in these studies

To test the effects of CF-MiPro on structure and function of a given gut microbiome sample, we utilized stool samples and colonoscopy aspirates (the latter harvested from the descending colon) in the passaging experiments outlined below. These clinical samples originate from nonCF donors (n=13) and CF donors (n=9). Clinical sample information is detailed in **Table 2**. In the experiments below, we compare the pooled stool/colonoscopy data from pwCF versus nonCF controls.

**Table 2.**
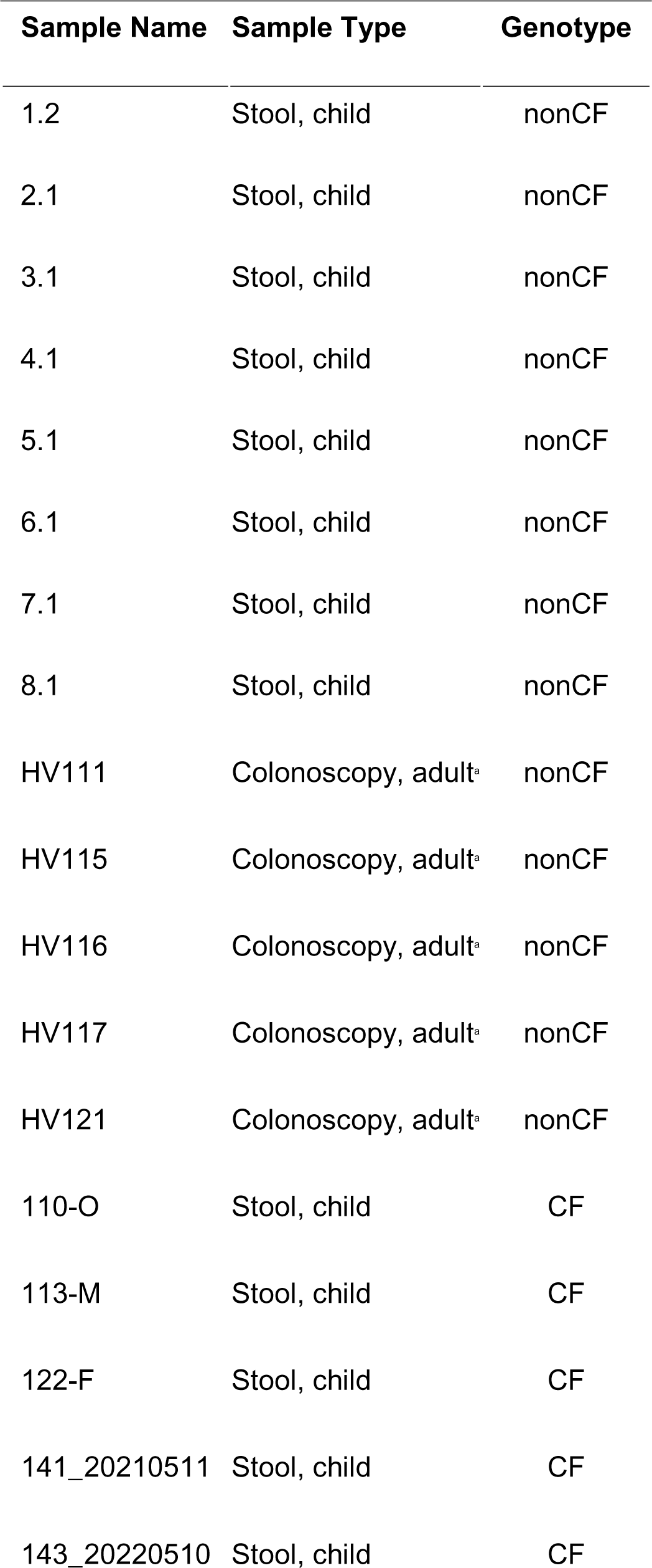
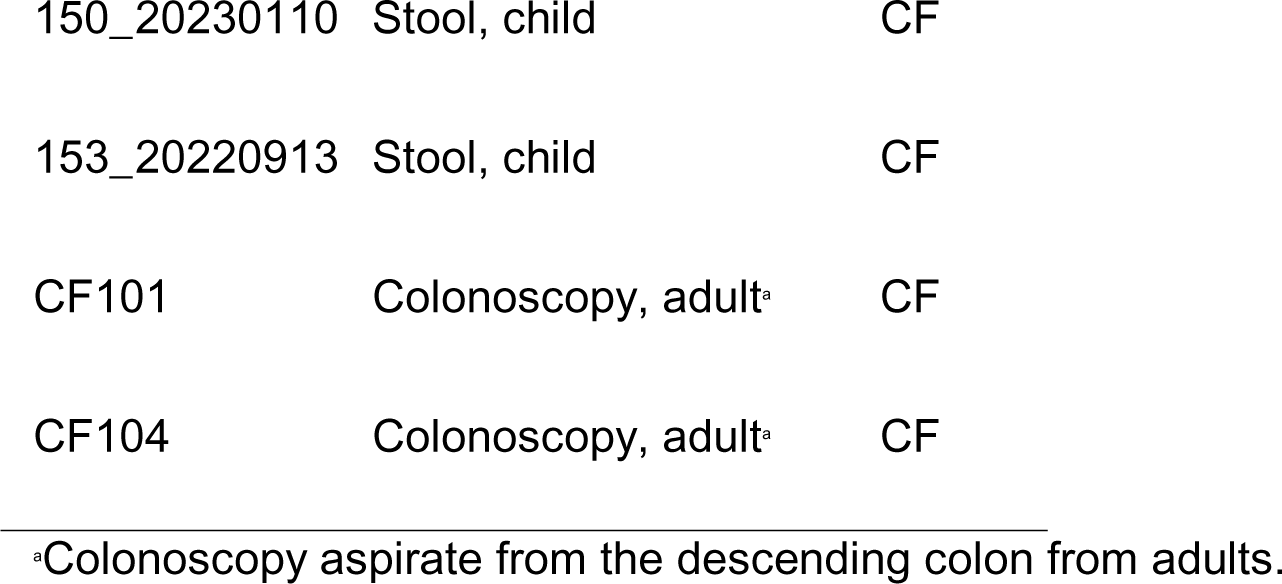
Source of stool/colonoscopy samples used in this study.

We determined the composition of these clinical samples using 16S rRNA gene amplicon library sequencing (**Table S1-2**). Overall, we note a non-significant difference in average Shannon Diversity Index (SDI) values for stool/colonoscopy samples from pwCF and nonCF genotypes, as analyzed by linear regression (**Figure S1A**; P=0.823). SDI is an alpha diversity measure encompassing both microbial richness and evenness. In agreement with these results, we found a non-significant difference in Chao-1 index, a measure of microbial richness, for stool/colonoscopy samples from pwCF and nonCF controls (**Figure S1B**, P=0.868).

The Bray-Curtis distance metric was used to determine the influence of genotype on microbial community similarity between samples and significance was tested with PERMANOVA. While there was modest overlap between these clinical samples from CF and nonCF sources, genotype played a significant role in beta diversity (**Figure S1C**, P=0.001). Interestingly, there are two nonCF outliers - these samples are both colonoscopy aspirates from adult donors. We do not know whether these samples were collected at a time of inflammation or illness.

We next calculated colony forming units per milliliter (CFU/mL) of each stool or colonoscopy aspirate following growth on blood sheep agar at 0% oxygen or 21% oxygen. Total CFU/mL is the sum of CFU/mL detected +/- oxygen (**Figure S1D**). Of note, total CFU/mL exaggerates the number of viable bacteria since some microbes will grow +/- oxygen (i.e., facultative anaerobes). We used a linear regression model to test whether genotype had a significant effect on cell count. CF samples cultured significantly less anaerobic CFU/mL (**Figure S1E**; P=0.0014) and aerobic CFU/mL (**Figure S1F**; P=0.045); thus, total CFU/mL was decreased in CF samples (**Figure S1D**; P=0.0017).

Lastly, we assessed broad changes in relative abundance at the phylum level between the two genotypes (**Figure S1G, Table S3**). When we analyzed the clinical samples based on origin, average Proteobacteria relative abundances were 14.5% higher in CF versus nonCF samples, which is in concordance with the literature (5, 15, 19). Also, Bacteroidota (Bacteroidetes) are decreased by 14.7% in CF samples compared to nonCF samples, also consistent with the literature (5, 15, 19). Actinobacteriota (nonCF: 14%, CF: 15.6%), Firmicutes (nonCF: 47.6%, CF: 46.2%) and Verrucomicrobiota (nonCF: 0.213%, CF: 0.333%) were not significantly different between the two genotypes (**Table S3**).

### CF-MiPro influences alpha and beta diversity following culture in vitro

To assess the impact of the media types tested here when culturing the stool and colonoscopy samples, homogenized stool or raw aspirates were inoculated at a 2% final volume/volume (v/v) ratio into MiPro, low-CF-MiPro or median-CF-MiPro – these initial cultures are designated “Day 0”. Each culture was then serially passaged anaerobically for five days. Each day, the cultures were homogenized by mixing, subcultured into fresh medium at 2% final inoculum, then incubated at 37°C for 24h before analysis. Each day we performed the following analysis: (i) CFU/mL were calculated by plating aliquots of the homogenized, planktonic culture on sheep blood agar that were then incubated at 0% or 21% oxygen, (ii) the cell pellets were collected for DNA extraction and 16S rRNA gene amplicon library sequencing, and (iii) culture supernatants were filter-sterilized and stored for subsequent analysis of short chain fatty acids (SCFA, **Figure 1A**).

**Figure 1.**
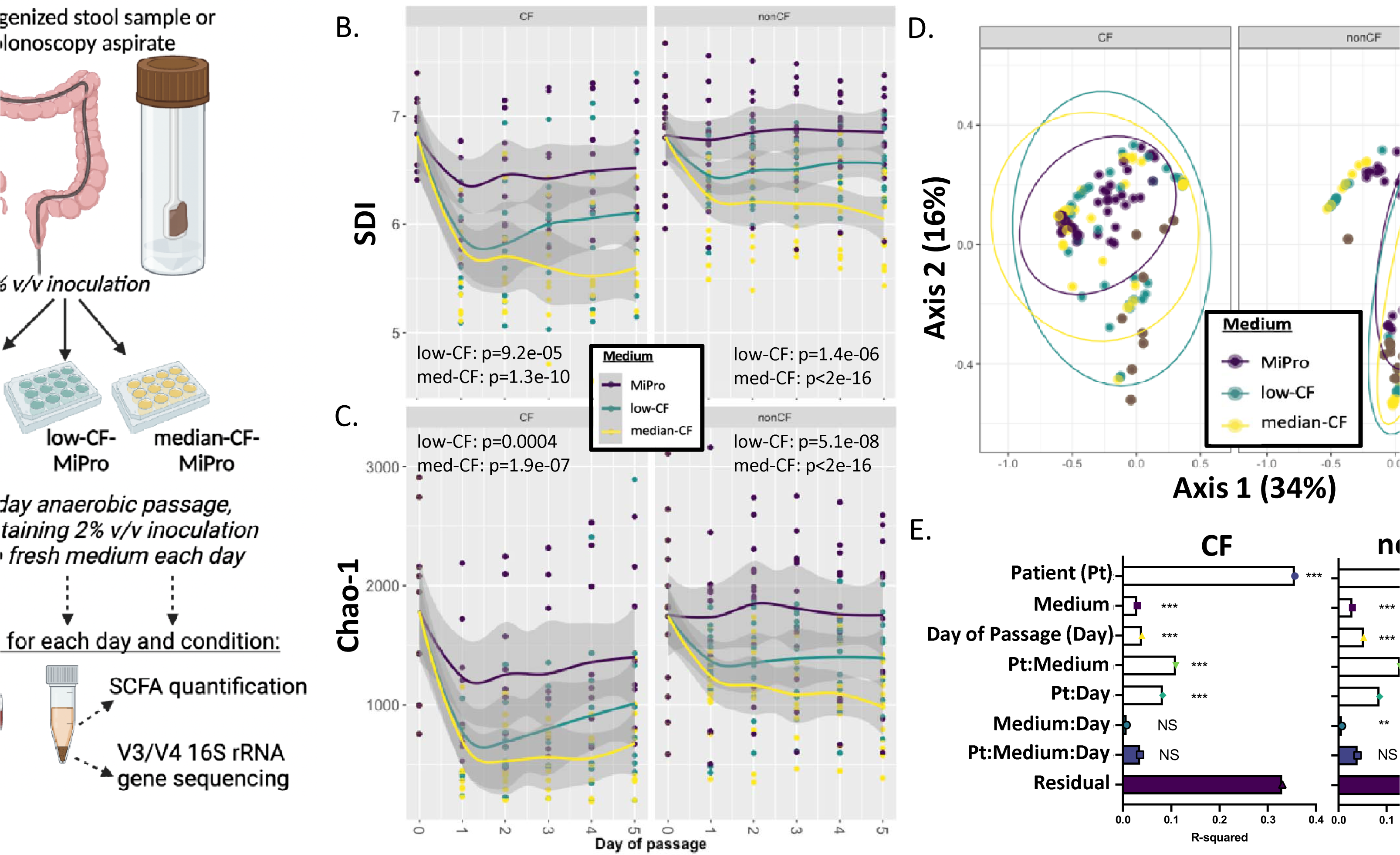
Impact of CF-MiPro on the diversity of cultured clinical samples from CF and nonCF subjects. **(A)** Schematic of the experimental design. Stool and colonoscopy samples are grown in the indicated medium over 5 days under anoxic conditions, then subjected to analysis as follows: viable counting, amplicon sequencing and targeted analysis of SCFA. **(B)** Shannon Diversity Index (SDI) and **(C)** Chao-1 distance of CF (left) and nonCF (right) samples cultured in MiPro (purple), low-CF-MiPro (teal) and median-CF-MiPro (yellow) for five days. Each day, a 2% inoculum was introduced into fresh medium, with the passages every 24h. Samples were collected at day zero and daily though the end of day 5. A linear mixed effect model from the R package lme4 was used to test whether SDI or Chao-1 changed significantly with medium type. Patient was set as the random variable to control for multiple sampling. Both SDI and Chao-1 are significantly negatively correlated with low- and median-CF-MiPro for both genotypes, compared to MiPro. **(D)** Bray-Curtis beta diversity was calculated for each sample and displayed on a principal coordinate analysis (PCA) plot, faceted by genotype and colored by medium. The first two components account for 50.2% of total variance. **(E)** Significant differences in beta diversity were tested by PERMANOVA with metadata included in the model, and R-squared values for all metadata included in model, as well as potential interactions, are plotted with significance codes indicated (NS: non-significant, **: p<0.01, ***: p<0.001). “Residual” indicates variation unexplained by the model.

We generated 16S rRNA gene amplicon library sequencing data from DNA extracted from all cell pellets from all passages. We analyzed these data using a mixed linear model for each genotype with patient (or sample source) as the random effect to test whether SDI was significantly correlated with medium and day of passage. For samples originating from a CF donor, medium and day of passage significantly correlated with decreased SDI in the CF-MiPro media formulations (**Figure 1B, left**; low-CF-MiPro, P=9.2e-05; median-CF-MiPro, P=1.3e-10; Day P=0.0002). The same trend is observed for samples originating from a nonCF donor, whereby medium significantly correlated with decreased SDI in a dose-dependent manner (**Figure 1B, right**; low-CF-MiPro, P=1.4e-06; median-CF-MiPro, P<2e-16; Day P=0.014).

A second metric of alpha diversity, Chao-1, which estimates total richness, indicated that both formulations of CF-MiPro reduced the number of taxa in a community across stool/colonoscopy samples from pwCF (**Figure 1C, left**; CF: low-CF-MiPro, P=0.0004; median-CF-MiPro, P=1.9e-07; Day P=0.0006) and from nonCF clinical samples (**Figure 1C, right**; nonCF: low-CF-MiPro, P=5.1e-08; median-CF-MiPro, P<2e-16; Day P=0.001).

To confirm the use of MiPro as a maintenance medium (33), we ran a mixed linear model for each genotype with patient as the random effect to test whether SDI and/or Chao-1 were significantly correlated with day of passage. Overall, MiPro maintains SDI (**Table S4,** CF: Day P=0.293; **Table S5,** nonCF: Day P=0.411) and Chao-1 (**Table S6**, CF: Day P=0.249; **Table S7,** nonCF: Day P=0.875) measures of a given inoculum after five days of in vitro passage. Thus, our analysis replicates the findings of Li and colleagues (33) with respect to MiPro as a maintenance medium.

The Bray-Curtis distance metric was then applied to the 16S rRNA gene amplicon library sequencing data and used to test the influence of patient, medium, day of passage, and their interaction terms on overall microbiome composition within each genotype (**Figure 1D**). Significance was tested by PERMANOVA (**Figure 1E**). Patient, medium and day of passage all significantly impacted community composition, with patient having the largest R^2^ value.

We also used Jaccard and Morisita-Horn as alternative metrics of beta diversity. Jaccard index measures the similarity between groups based on occurrence measures (i.e., presence or absence of taxa) and does not consider relative abundances of taxa. Morisita-Horn index accounts for relative abundance of taxa but ignores taxa with 0% relative abundance: this metric only clusters by presence of shared taxa, rather than both presence and absence. PCA plots displaying both Jaccard index (**Figure S2A**) and Morisita-Horn index (**Figure S2B**) resemble each other, and the Bray-Curtis distance (**Figure 1D**). The first two components of the PCA plot displaying Jaccard index account for 36.9% of total variance, and 60% of total variance is displayed by Morisita-Horn. For each beta diversity metric, patient contributes to the most variance for both CF and nonCF samples, followed by medium and day of passage, which contribute to similar variance between both genotypes (**Figure S2C-D**). Residual values for Jaccard analysis are larger than those for Morisita-Horn, which may support the robustness of accounting for both compositionality and absent taxa.

It is notable that all combined exposures included in the model explained only 67-73% of the total variance in beta diversity in CF and nonCF samples, respectively. The residual variance indicates that there is a modest amount of variation between samples that is not due to any of the factors examined here. To address this residual variance, we incorporated sample preparation (“Prep”) as an explanatory variable. Because four CF samples were initially homogenized in PBS+10 mL L-cysteine, rather than PBS+7.15% glycerol, we calculated the Bray-Curtis distance metric on all CF samples at Day 5 of passage (**Figure S3A**). Here, we aimed to address the role of different preparation methods on final microbial composition following five days of in vitro passage. Principal components analysis (PCA) plots displaying Bray-Curtis beta diversity suggest that samples prepared in L-cysteine are more similar to each other, and samples prepared in glycerol are more variable. Importantly, cysteine-prepared samples seem to maintain microbial composition across media conditions. The first two components of the PCA plot account for 61.1% of total variance. Patient contributes to the most variance in CF Day 5 samples (**Figure S3B**, R^2^=0.418, P=0.001), followed by preparation (R^2^=0.133, P=0.003), then medium (R^2^=0.0715, P=0.163). These data suggest the significant role of preparation on final microbial composition and more so than growth in CF-MiPro. Based on these findings, PBS+10 mL L-cysteine should be used to prepare the samples in future studies.

We note four important findings from the analysis above. First, both CF and nonCF genotype significantly affect how community composition changed over days of passage (Patient:Day of passage), indicating differential rates of change within individuals, a finding consistent with the observation that “patients are most like themselves” when analyzing microbiota. Second, for nonCF samples, CF-MiPro medium significantly affected how community composition changed over days of passage (Medium:Day of passage), suggesting continuous shifts in microbial composition as the microbiota adapted to the CF-MiPro medium conditions. Third, for measures of α- and β-diversity, for both CF and nonCF samples, there was a significant interaction between patient and medium, supporting the distinct role of each individual’s response to the CF-MiPro medium. Finally, because the interaction term “Medium:Day of passage” is non-significant in altering CF community composition, we can infer that shifts in beta diversity within media types are largely static across days for this genotype. We highlight these latter two important points in the Discussion. In conclusion, these results suggest that the patient contributes the most to differences in beta diversity (independent of analytical tool used), followed by medium, method of preparation and day of passage. Notably, the microbial composition of CF samples remain relatively static over the course of passaging, while nonCF microbial compositions continue to shift over time in response to growth in CF-MiPro medium.

### CF-MiPro induces changes in relative abundance of microbiota over time to a more CF-like state

To understand the general shifts that occur across in vitro conditions, we first examined broad changes in relative abundance that occur over time at the phylum level. We compared average phylum-level relative abundances from all samples collected over the five-day passage within each medium and genotype (**Figure 2A**, **Table S3**). In general, phylum-level relative abundances in passaged samples from pwCF remain stable over five days within all three media types (**Figure 2A, top**), although there are modest shifts in average relative abundance following in vitro passage in MiPro and low-CF-MiPro. For example, Firmicutes increase slightly from an average of 63% in raw CF samples to 74.4% over the course of in vitro passage in MiPro, ∼60% in low-CF-MiPro and ∼72% in median CF-MiPro (**Table S3**). Additionally, Actinobacteria decrease from 12.3% in raw CF samples to ∼1.5% in all media formulations. These shifts likely occur due to the brief introduction of oxygen during sample preparation and acclimation to an in vitro system. The average relative abundances of the other top phyla remain generally stable between cultivation in MiPro and median-CF-MiPro, indicating that CF-MiPro can serve as a maintenance medium for CF gut microbiome samples (**Table S3**).

**Figure 2.**
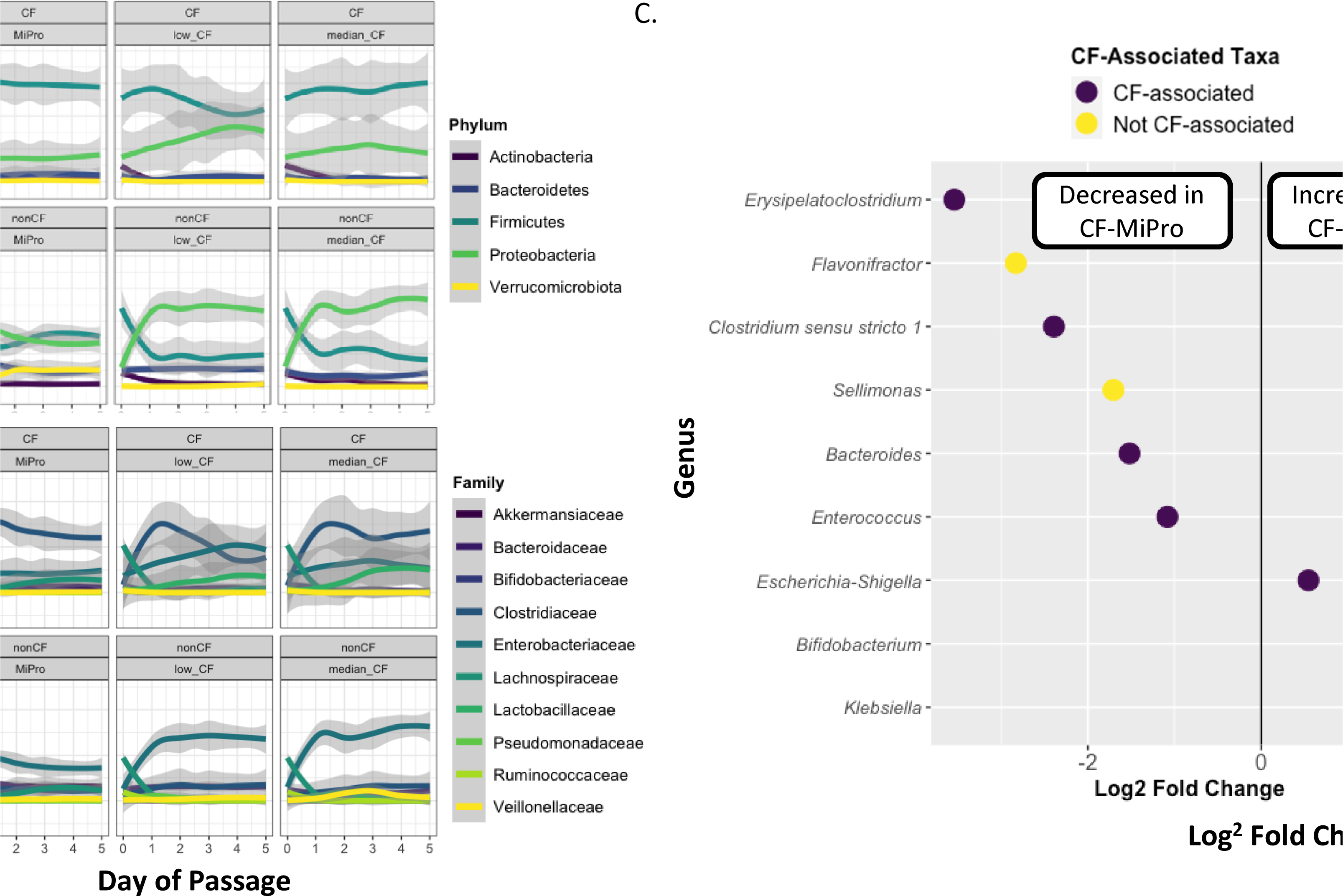
Microbial relative abundance changes across media types. Day of passage is graphed versus relative abundance of the indicated taxa for each sample, and a ribbon plot was used to visualize overall changes in microbial relative abundance at the (**A**) phylum level and (**B**) family level across all media conditions. The mean relative abundance is depicted by the line with 95% confidence interval shaded in grey. The legend indicates the taxonomic assignment for each panel. (**C**) Log2 fold change of taxa that were significantly (p-adj.<0.05) altered in nonCF samples passaged in median-CF-MiPro versus MiPro. Taxa were filtered to be present in at least 5% of all samples. Each dot represents a single genus and is color coded by association with CF gut dysbiosis (see main text for details). Significance was determined by DESeq2 using a non-continuous model of samples binned by media condition across days of passage 1 through 5. Patient was included as a design variable to control for multiple sampling.

Phylum-level relative abundances in nonCF samples differ significantly from CF samples in terms of shifts in microbiota across the different media types. Shifts observed in MiPro occur at Day 1 of passage in a batch-independent manner (**Figure S4**), suggesting the sensitivity of nonCF gut microbiota to oxygen and/or cultivation in this medium likely due to the significantly larger concentration of detectable anaerobic microbes in nonCF stool (**Figure S1E**). The largest overall changes occur in (i) Firmicutes, which decreases from an average of 62.3% in raw nonCF samples to 36.1% over the course of passage in MiPro, and (ii) Proteobacteria, which increases from an average of 12.3% in raw nonCF samples to 36.9% over the course of passage in MiPro (**Figure 2A, bottom; Table S3**). Importantly, these changes we observe in nonCF samples in MiPro are exacerbated in a dose-dependent manner in low- and median-CF-MiPro over five days of passage. For example, Firmicutes average relative abundance further decreases from 36.1% in MiPro to 23.7% in median-CF-MiPro, while Proteobacteria increase from 36.9% in MiPro to 64.7% in median-CF-MiPro (**Table S3**). Bacteroidota and Verrucomicrobiota relative abundances also change unidirectionally, with Bacteroidota decreased from 13.5% in MiPro passages to 8.32% in median-CF-MiPro passages, and Verrucomicrobiota reduced from 10.8% in MiPro to 0.0828% in median-CF-MiPro passages (**Table S3**). These changes are consistent with comparisons of persons with CF with nonCF cohorts (5, 15, 18, 19).

Our analysis show that in the clinical samples used here, the phyla Actinobacteria, Proteobacteria, Bacteroidota and Verrucomicrobiota are dominated by single families: Bifidobacteriaceae, Enterobacteriaceae, Bacteroidaceae and Akkermansiaceae, respectively. Phylum-level changes appear to be driven primarily by changes in these taxa (**Figure 2B, Table S8**). In nonCF raw samples, Lachnospiraceae appears to be the dominant Firmicutes family (39.2% relative abundance, **Table S8**). In CF raw samples, Lachnospiraceae is also the dominant Firmicutes family, with a relative abundance of 42.9%. However, when passaged in CF-MiPro, Lachnospiraceae is decreased and Clostridiaceae dominates for both genotypes. These observations align with literature reporting the increased relative abundance of Clostridiaceae (48) and reduced Lachnospiraceae in CF stool (14).

We next determined the genera that changed significantly between media conditions across all samples. We first compared average genus-level relative abundances from all samples collected over the five-day passage within each medium and genotype (**Figure S5, Table S9**). The families Bifidobacteriaceae, Clostridiaceae, Enterobacteriaceae, Bacteroidaceae and Akkermansiaceae are dominated by single genera: *Bifidobacteria*, *Clostridium*, *Escherichia/Shigella*, *Bacteroides* and *Akkermansia*, respectively. Family-level changes appear to be driven primarily by changes in these taxa (**Table S9**). For passaged samples from pwCF, *Clostridium* is the dominant genus within the Firmicutes phylum in MiPro and CF-MiPro. In nonCF samples, *Blautia* is the dominant genus in raw samples yet is replaced by *Clostridium* following passage in CF-MiPro.

Next, we wanted to determine which genera may be the most significantly enriched or depleted in CF-MiPro compared to MiPro. To perform this analysis, we compared a subset of samples within each genotype grouped by medium: samples cultured in MiPro and samples cultured in median-CF-MiPro. We excluded raw samples from this analysis. We then filtered by taxa that are present at least 5% relative abundance of all samples (prevalence>0.05) and those with relative abundances that significantly change between MiPro and median-CF-MiPro (p-adj.<0.05 by Wald test statistic). Taxa that changed significantly across these conditions were compared with taxa associated with CF gut microbial dysbiosis (51). Among the 9 genera that shifted significantly between media types for nonCF-passaged samples, 7 genera were associated with CF gut dysbiosis. For example, *Escherichia*, specifically *E. coli*, is reported to be enriched in CF stool (5, 15, 18, 19). We found that *Escherichia/Shigella* is significantly enriched across samples passaged in median-CF-MiPro, compared to MiPro (log2FC=0.54, p-adj.=0.02; **Figure 2C**). Additionally, SCFA-producing microbes, such as *Bacteroides* and *Akkermansia*, are reported to be depleted in pwCF. Similarly, we see a 1.52-fold decrease in *Bacteroides* in median-CF-MiPro (p-adj.=2.2E-11). Of note, *Akkermansia* decreases by 1.04-fold in median-CF-MiPro among nonCF samples, however this difference is non-significant compared to passage in MiPro (p-adj.=0.52, **Table S10**).

Interestingly, we observed a significant decrease in two *Clostridium*-related taxa (*Erysipelatoclostridium* and *Clostridium sensu stricto 1*). There have been mixed reports of *Clostridium* relative abundances in CF (14, 21, 48–50), some of which are associated with antibiotic use (21). These results suggest intra-genus diversity among pwCF and the putative association of certain species with infection and antibiotic perturbation. Because our study lacked metadata for each patient and we used 16S rRNA gene sequencing, further investigation is needed to discern which species are depleted in median-CF-MiPro and if these species are correlated with clinical outcomes. Of note, when we performed this analysis on a subset of CF samples passaged in MiPro and median-CF-MiPro, we were not able to identify any genera that shifted in relative abundance under the same parameters used in the nonCF analysis (prevalence>5%, p-adj.<0.05). These results support that CF-MiPro maintains CF gut microbiome samples at the genus level, yet significantly shifts nonCF gut microbiome samples toward a “CF-like” microbial profile with less *Bacteroides* and more *Escherichia*.

Overall, these results highlight: (i) the highest observed change in taxa abundance occurs when nonCF samples are passaged in CF-MiPro medium, and these changes are larger for median-CF-MiPro versus low-CF-MiPro, indicating a “dose-dependent” response, (ii) the overall patterns of change appear to be driven by a small number of bacterial families, and often a sole family, (iii) the shifts observed in CF-MiPro largely align with clinical observations of the shifts in phyla and genera associated with pwCF, and importantly, (iv) for nonCF samples, passaging in CF-MiPro results in a shift in these samples to a CF-like microbiota composition.

### CF-MiPro decreases absolute microbial abundance

We next examined CFU/mL of each sample as a read-out of the culturable, absolute abundance of the microbial population. Inocula (raw samples) and in vitro passaged cultures were serially diluted 10-fold and plated on blood sheep agar, then incubated at either 0% or 21% oxygen for 24h. We used a mixed effect linear model for each genotype with patient as the random effect to test whether CFU/mL was significantly correlated with medium or day of passage. As previously described, raw CF samples cultured less aerobic and anaerobic CFU/mL (**Figure S1D-F**). In conjunction with decreased richness observed in samples cultured in CF-MiPro (**Figure 1C**), it was our expectation that CF-MiPro would foster fewer total microbes. Across clinical samples from both genotypes cultured through MiPro, low-CF-MiPro and median-CF-MiPro for five days, CF-MiPro showed decreased anaerobic CFU/mL in a dose-dependent manner (**Figure 3A**). This decrease in anaerobic CFU/mL in CF-MiPro is day-dependent in nonCF cultures (Day P=0.00018) and in CF cultures (Day P=0.045). Similarly, the aerobic CFU/mL fraction is reduced over passages in all media; although, this reduction is observed in a day-independent manner for both genotypes and the reductions are more modest (**Figure 3B**).

**Figure 3.**
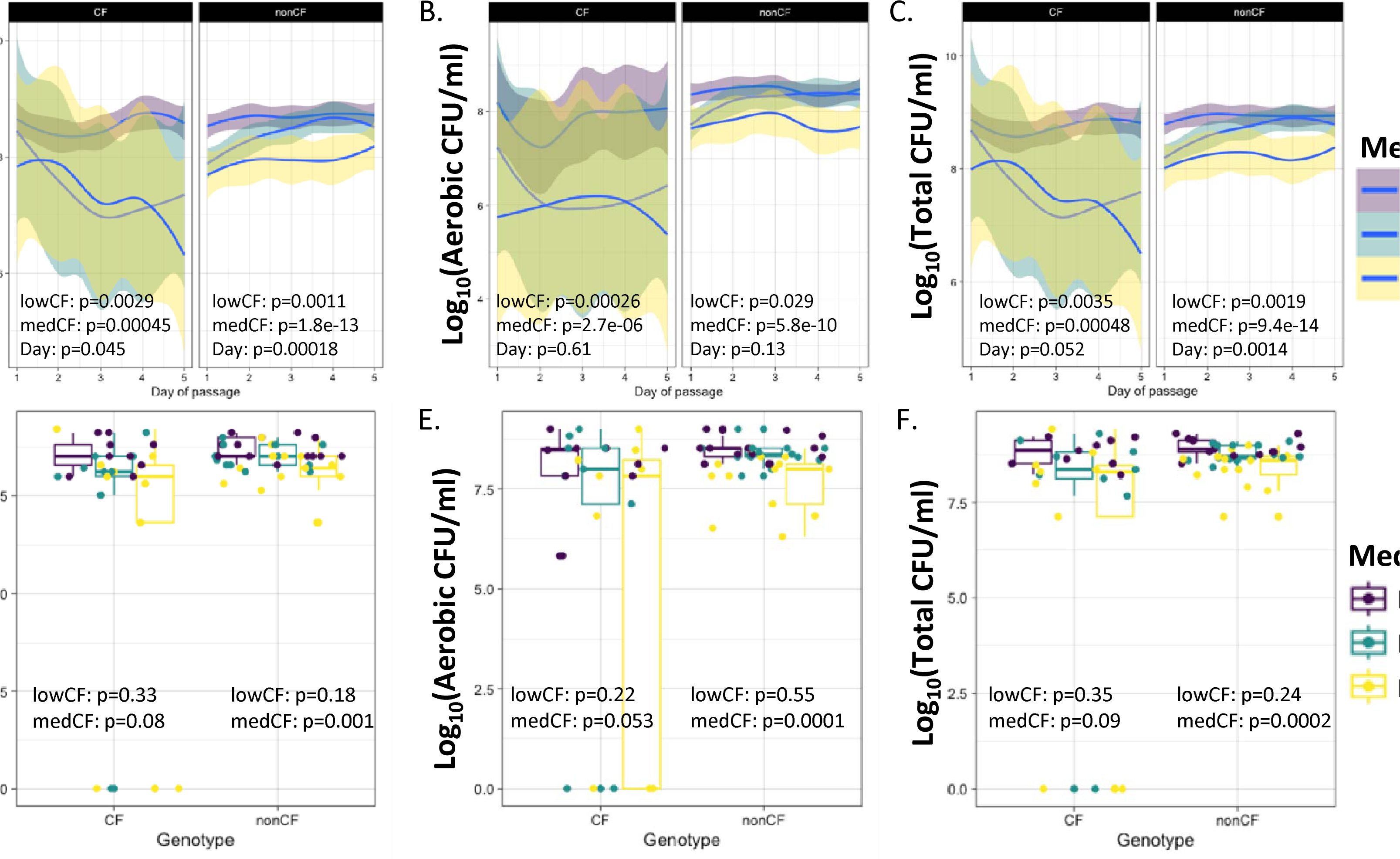
CFU/mL of nonCF and CF stool or colonoscopy samples passaged in CF-MiPro over five days. NonCF (n=13) and CF (n=9) stool and colonoscopy samples were inoculated into each medium (MiPro, purple; low-CF-MiPro, teal; median-CF-MiPro, yellow) at a 2% final v/v ratio and passaged for 5 days, using this 2% inoculum into fresh medium daily. Each day, viable cell counts were quantified on sheep blood agar plates incubated at (**A**) 0% oxygen for obligate anaerobe and facultative anaerobe detection or (**B**) 21% oxygen for aerobic/facultative anaerobe detection. Total CFU/mL (log10(aerobic+anaerobic CFU/mL)) are displayed in (**C**). A linear mixed effect model from the R package lme4 was used to test whether CFU/mL within each genotype changed significantly with medium type and day of passage. Patient was set as the random variable to control for multiple sampling. CFU/mL are typically negatively correlated with CF-MiPro, in a dose-dependent manner, for the cultured CF and nonCF stool and colonoscopy samples, suggesting a decrease in richness. Overall, day of passage is positively correlated with CFU/mL for CF and nonCF cultured samples. To eliminate the contribution of Day of Passage, (**D**) anaerobic, (**E**) aerobic and (**F**) total CFU/mL from Day 5 are graphed by genotype, colored by medium. Another linear mixed effect model, with patient set as the random variable, was used to test whether Day 5 CFU/mL within each genotype changed significantly with medium. Under all oxygen tensions, CFU/mL from CF samples used as inoculum were not correlated with medium, while CFU/mL from nonCF samples used as inoculum were negatively correlated with median-CF-MiPro, but not low-CF-MiPro.

CF-MiPro is significantly correlated with decreased total CFU/mL (sum of aerobic and anaerobic CFU, then log10-transformed) for clinical samples from both subject genotypes, in a day-dependent manner for nonCF samples (Day, P=0.0014) and marginally significant in CF samples (Day: P=0.052). To account for day-to-day variability, we next graphed CFU/mL calculated from Day of Passage 5 from each oxygen tension. Linear regression, with medium as the explanatory variable, was used for statistical analysis. Anaerobic CFU/mL at Day 5 are not significantly affected by medium in cultures inoculated with CF samples (**Figure 3D, left**; low-CF-MiPro, P=0.33; median-CF-MiPro P=0.08), but median-CF-MiPro significantly decreases Day 5 anaerobic CFU/mL in cultures inoculated with nonCF samples (**Figure 3D, right**; low-CF-MiPro, P=0.18; median-CF-MiPro, P=0.001). This observation is repeated for the aerobic Day 5 CFU/mL, whereby only cultures inoculated with nonCF samples are decreased in median-CF-MiPro (**Figure 3E**). Decreased total CFU/mL in cultures inoculated with nonCF samples, but not CF samples, are correlated with median-CF-MiPro (**Figure 3F**). Taken together, these results suggest that samples from both genotypes are affected by CF-MiPro, as evident by the trend of decreasing CFU/mL in low- and median-CF-MiPro, but only nonCF samples are statistically-significant, likely due to the greater variability and more modest reductions for CF samples.

### CF-MiPro shifts short chain fatty acid profiles

Short chain fatty acids are metabolic byproducts of dietary fiber catabolism (51). These metabolites are the preferred nutritional source for host enterocytes and are known to have antimicrobial and anti-inflammatory properties (52). Two studies have reported a significant reduction in SCFA concentrations in CF stool compared to nonCF stool (48, 50), an observation that is likely due to the reduction in SCFA-producing microbes (53) and/or the enrichment of KEGG-annotated SCFA catabolism genes in CF stool (16). It was our prediction that, given the shift to a more CF-like microbiota in CF-MiPro, passages in this medium would lead to a decrease in SCFA concentrations over the course of five days of passaging.

To analyze the impact of medium on SCFA production, filtered supernatants from all cultured samples (nonCF, n=8; CF, n=5) collected through Day 1-5 were analyzed using gas chromatography-mass spectrometry (GC/MS). SCFA concentrations were determined by normalization to standards, with a detection limit of 0.3 uM. Then, these concentrations were normalized by baseline levels detected in each respective medium (MiPro, low-CF-MiPro and median-CF-MiPro). Finally, these concentrations were normalized to the CFU/mL calculated in that exact sample to account for differences in microbial density. We used a mixed linear model for each genotype with patient as the random effect to test whether each SCFA measured (acetate, propionate, butyrate) was significantly correlated with medium or day of passage.

For samples from pwCF, acetate is not significantly correlated with either medium or day (**Figure 4A, left**). However, butyrate and propionate concentrations are reduced in CF-MiPro, in a dose-dependent manner (**Figure 4A, middle and right**). In nonCF samples, all three SCFA concentrations are negatively correlated with CF-MiPro, with acetate and propionate concentrations positively correlated with day (**Figure 4B**).

**Figure 4.**
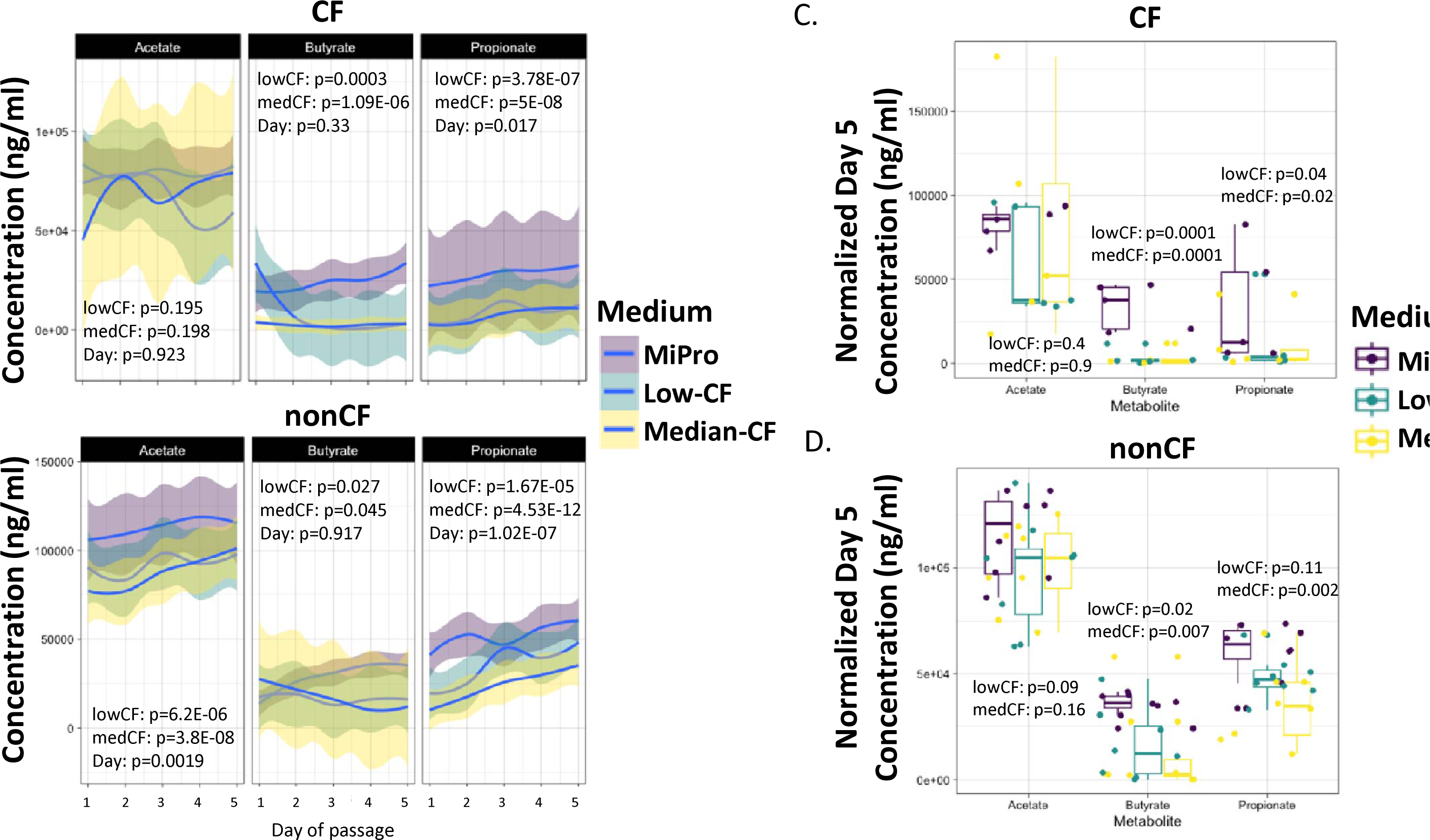
Short-chain fatty acid (SCFA) production is decreased in CF-MiPro. Filtered supernatants from Day 1-5 cultures were analyzed via GC/MS for quantification of the three most abundant SCFA: acetate, butyrate and propionate. The final concentrations were normalized to total anaerobic CFU/mL and graphed against Day of Passage for (**A**) CF and (**B**) nonCF samples. A linear mixed effect model from the R package lme4 was used to test whether each SCFA concentration within each genotype changed significantly with medium type and day of passage. Patient was set as the random variable to control for multiple sampling. While acetate concentrations were unaffected, butyrate and propionate concentrations were negatively correlated with CF-MiPro when CF samples were used as inoculum. In contrast, all three SCFA were negatively correlated with CF-MiPro when nonCF samples were used as inoculum. Day of passage inconsistently had a positive correlation with SCFA concentrations. To eliminate the contribution of Day of Passage, concentrations of each SCFA from Day 5 are plotted for (**C**) CF samples and (**D**) nonCF samples. Statistical analysis was performed using a similar linear mixed effect model as in (**A**) and (**B**). At Day 5, butyrate and propionate concentrations are negatively correlated with CF-MiPro across both genotypes of clinical samples.

To account for day-to-day variability, we next graphed SCFA concentrations from Day of Passage 5 for each genotype. Linear regression, with medium as the explanatory variable, was used for statistical analysis. In cultures inoculated with CF samples, acetate concentrations remain unaffected by medium, and butyrate and propionate concentrations are both significantly reduced in low- and median-CF-MiPro (**Figure 4C**). Similarly, acetate concentrations are unaffected by medium for nonCF samples as the inoculum, while butyrate and propionate concentrations are both significantly reduced in CF-MiPro, in a dose-dependent manner (**Figure 4D**).

We next examined how overall changes to the SCFA levels, including normalized concentrations of acetate, butyrate and propionate, changed across media conditions at Day 5 of passage. Here, we used the Bray-Curtis distance metric to test the influence of patient and medium on overall SCFA composition in CF samples (**Figure 5A**) and nonCF samples (**Figure 5B**). Significance was tested by PERMANOVA within each genotype.

**Figure 5.**
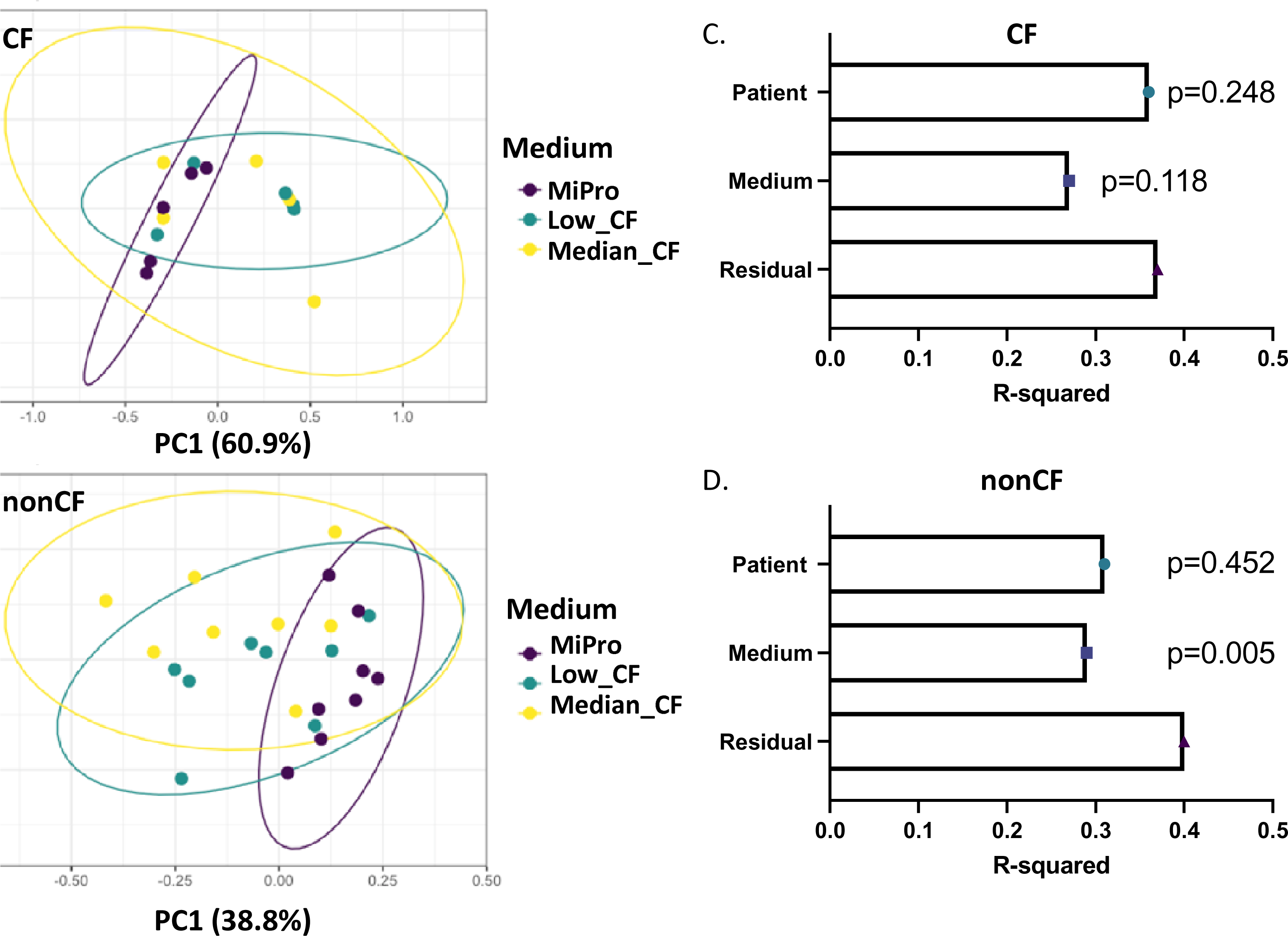
Short-chain fatty acid (SCFA) profiles are distinguished by medium in nonCF samples. CFU-normalized concentrations of acetate, butyrate and propionate at Day 5 of passage were used to calculate Bray-Curtis beta diversity for (**A**) CF and (**B**) nonCF stool and colonoscopy samples. Ordination of each genotype was displayed in principal coordinate analysis (PCA) plots, colored by medium. For CF-derived samples, the first two components account for 72% of total variance. For nonCF-derived samples, the first two components account for 53% of total variance. Significant differences in beta diversity for (**C**) CF and (**D**) nonCF samples were tested by PERMANOVA with metadata included in the model, and R-squared values for Medium and Patient are plotted with respective p-values. “Residual” indicates variation unexplained by the model. SCFA profiles in nonCF samples are significantly distinguished by medium in a patient-independent manner, whereas SCFA profiles in CF samples are not distinguishable by medium.

For samples from pwCF as the inoculum, the first two components of the PCA plot accounted for 71.92% of total variance (**Figure 5A**), with patient having the largest R^2^ value, with a nonsignificant correlation with beta diversity (**Figure 5C**, R^2^=0.36, P=0.248). Similarly, medium non-significantly correlated with beta diversity (**Figure 5C**, R^2^=0.27, P=0.118) for samples from pwCF.

For nonCF samples used as the inoculum, the first two components of the PCA plot accounted for 52.54% of total variance (**Figure 5B**), with patient having the largest R^2^ value, with a nonsignificant correlation with beta diversity (**Figure 5D**, R^2^=0.31, P=0.452). NonCF SCFA profiles are significantly influenced by medium, in a patient-independent manner (**Figure 5D**, R^2^=0.29, P=0.005). Both exposures included in the model (patient and medium) explained only 60-63% of the total variance in beta diversity in nonCF and CF samples, respectively. The residual variance indicates that there is a modest amount of variation between samples that is not due to any of the factors examined here.

Overall, these results suggest that CF-MiPro is significantly correlated with reduced levels of SCFA, with a larger effect size when nonCF samples are used as inoculum versus samples from pwCF. In contrast, for MiPro, there is a non-significant change in SCFA regardless of inoculum, consistent with the reports that MiPro can maintain gut microbiota from clinical sources in vitro.

## Discussion

In this study, we developed a novel first-generation in vitro culture medium, called CF-MiPro, to mimic the physiological conditions of the CF colon. Similar to the early approaches used in the generation of Artificial Sputum Medium (ASM; (54, 55)), a culture medium used to mimic the nutritional conditions of the CF airway, we incorporated validation of CF-MiPro as part of our analysis. We adopted a previously used experimental approach (33) to track community structure (utilizing 16S rRNA gene amplicon sequencing) and function (SCFA quantification) of 22 gut microbiome samples over time for these media.

Firstly, our results highlight that the composition of CF gut microbiome samples is generally maintained in CF-MiPro. In low- and median-CF-MiPro we observe similar trends, whereby CF gut microbiome samples stabilize with similar relative abundances of the top five phyla. In fact, no genera were identified to be significantly enriched or depleted in median-CF-MiPro when stool or colonoscopy samples from pwCF were passaged. Beta-diversity analysis of community composition results in a non-significant interaction between medium and day of passage in CF-derived samples, suggesting that day of passage does not alter microbial composition for CF-MiPro medium. Thus, community composition of CF samples remains relatively stable over the five days of passage when grown in CF-MiPro.

Conversely, nonCF gut microbiome samples undergo significant taxonomical shifts when introduced to CF-MiPro, in a dose-dependent manner. That is, passage of these nonCF stool or colonoscopy samples as inocula in low-CF-MiPro induces taxonomical shifts that are exacerbated in median-CF-MiPro. Our results highlight that the highest-abundance taxa change with medium, with overall patterns being driven by a small number of bacterial families and genera. These shifts observed in CF-MiPro generally align with clinical observations associated with pwCF. For example, analysis of stool from cwCF across cohorts, both nationally and globally, consistently report an increase in Proteobacteria, largely driven by *Escherichia coli* (5, 15, 18, 56). When nonCF samples are passaged in CF-MiPro, *E. coli* dominates the community after 24h. Over the course of a five-day passage, *Escherichia-Shigella* is significantly enriched in median-CF-MiPro. A key clinically-relevant microbe that is reported to be depleted in the CF gut, *Bacteroides*, is observed to be significantly depleted in median-CF-MiPro over the duration of in vitro passaging. These results suggest that, for nonCF samples, passaging in CF-MiPro promotes a shift to a CF-like microbiota composition. Furthermore, beta-diversity analysis of nonCF-derived samples as inocula result in a significant interaction between medium and day of passage, suggesting that day alters community composition within each medium. That is, the communities become more “CF-like” over time, as indicated through shifts in CF-associated taxa, and the gut microbiota of these nonCF samples is dynamically shifting over the five-day duration, thus presumably continuously responding to one or more components in CF-MiPro.

For both CF and nonCF samples cultivated in each medium type, we report a significant interaction between patient and medium. This result underscores the important role of person-to-person variation in response to different stimuli, in this case medium. Furthermore, in every analysis, patient is reported to have the largest R-squared value, indicating the largest contribution to variance. These results emphasize the need to account for systematic differences between patients in all statistical approaches. Here, we account for patients through mixed effect models by setting patient as the random variable.

In our functional analysis, which serves as a second means to validate the utility of CF-MiPro, we tested SCFA concentrations in cultivated samples. We report a significant dose-dependent depletion of butyrate and propionate in CF-MiPro, compared to MiPro. This depletion is observed in both CF and nonCF samples and aligns with the reduction in SCFA-producing microbes, such as *Bacteroides*. However, when comparing the overall SCFA composition (inclusive of acetate, butyrate and propionate concentrations) at Day 5 via Bray-Curtis dissimilarity, medium has a significant effect on SCFA profiles in nonCF samples, but not CF samples. In fact, medium significantly contributes to nonCF SCFA concentrations in a patient-independent manner. These results support the overall maintenance of function for samples derived from pwCF in CF-MiPro, as tested through one functional output: SCFA quantification. As a control, we show that MiPro, as reported previously (33), maintains both the population composition and SCFA concentrations over the course of the experiment.

Overall, we view this first-generation CF-MiPro formulation as another tool, along with in vitro tissue culture/organoid and animal models, to better understand the microbiota dysbiosis associated with CF. Despite the ability of CF-MiPro medium to maintain compositional and functional features of CF microbial communities derived from the gut, we note a limitation of CF-MiPro: it uses non-physiological substrates in its formulation. The source of “fat” in CF-MiPro is glycerol; the actual in vivo source of fat is triglycerides and their breakdown products (15, 57). Similarly, the source of sugars is from yeast extract (i.e., mannose from the yeast cell wall, glucose and glycogen as intracellular carbohydrate sources) (33, 58) rather than the complex carbohydrates typically found in the gut (i.e., oligosaccharides and fibers) (59). Other features, like bile, are poorly characterized (or not investigated at all) in young CF cohorts. Future work will focus on modifying features of CF-MiPro as new data emerge describing the intestinal environment. This iterative process will help us to better reflect the CF intestinal environment in vitro and identify key features associated with microbial dysbiosis.

## Materials and Methods

### Samples, culture conditions and sequencing

A total of 22 samples (13 nonCF, 9 CF) were used for these experiments. Among the 13 nonCF samples, 8 were from a raw stool sample collected from children (approximate age <18-36 months), and 5 were from a colonoscopy aspirate from the descending colon of adults. Among the 9 CF samples, 7 were from a raw stool sample collected from children (approximate age <18-36 months), and 2 were from a colonoscopy aspirate from the descending colon of adults. See **Table 2** for details. The stool samples were thawed, weighed and resuspended in sterile PBS (Corning Cat. #21-040-CM) supplemented with 7.15% glycerol (and for some samples with L-cysteine added at 10 mM, see **Figure S2**) in a ratio of 1 to 5.7 w/v. The colonoscopy aspirates were frozen without processing, or without addition of glycerol.

For the passaging experiments, each sample was homogenized and inoculated into MiPro, low-CF-MiPro and median-CF-MiPro in a sterile 12-well plate (Corning Cat. #3512) to a final v/v ratio of 2% - this culture is designated “Day 0”. Each culture was passaged anaerobically for five consecutive days (24h per passage) in either a GasPak jar or an anaerobic chamber. Each day, the culture was homogenized by mixing and sub-cultured into fresh media at a 2% final v/v ratio. The remaining culture was analyzed as follows: (i) CFU/mL were calculated by plating aliquot of the planktonic culture on sheep blood agar that was then incubated at 0% or 21% oxygen, (ii) the cell pellets were collected and stored in QIAGEN RNAprotect (Cat.# 76506) for DNA extraction and 16S rRNA gene amplicon library sequencing. Homogenized stool samples or colonoscopy aspirates, as well as cell pellets from each day of passage, were stored at -80C and later processed with Zymo fecal DNA miniprep kit (Cat.# D6010). Paired-end reads were generated with 2x301 Illumina NextSeq2000 amplicon sequencing of the V3-V4 hypervariable region of the 16S rRNA gene (SeqCenter, LLC. Pittsburgh, PA). Raw data have been uploaded to the NCBI sequence read archive under BioProject PRJNA999361, and (iii) culture supernatants were filter-sterilized and stored at -80°C for subsequent analysis of short chain fatty acids (SCFA).

### Processing of rRNA-encoding gene amplicons

Primer sequences were removed by CUTADAPT (version 1.18). All subsequent preprocessing steps were performed in R version 4.2.2. The code is available at https://github.com/GeiselBiofilm/. A total of 61,916,801 raw paired-end reads were filtered and trimmed with DADA2 version 1.24.0. Reads were then denoised, merged, and chimeras removed. The final counts were 23,804,538 total reads, 60,112 mean reads per sample, and 42,103 unique amplicon sequence variants (ASVs). Taxonomy was assigned with DADA2 and the Silva version 138.1 training set. ASV taxonomy tables and sample metadata are available in supplemental tables (**Table S1** and **S2**). ASV counts for each sample are available at https://github.com/GeiselBiofilm.

### Analysis of rRNA-encoding gene amplicons

All downstream analysis and visualization were performed in R (version 4.2.2) Phyloseq (version 1.40) and ggplot2 (version 3.4.2) were used for data handling and visualization unless otherwise noted. Reads per sample were graphed and filtered to include only samples with >9,500 reads. All samples had at least 9,500 reads following processing and were, thus, included. Linear mixed models were used for statistical regression analysis (lme4 version 1.1.34). For each sample, beta diversity was calculated by Bray-Curtis, Morisita-Horn and Jaccard distances, and multidimensional scaling ordination was performed. Significant differences in beta diversity were tested by permutational analysis of variance (PERMANOVA) (vegan version 2.6.4).

### Metabolite quantification and analysis

Undiluted, filter-sterilized supernatants from Day 1-5 cultures were stored frozen at -80C prior to metabolite quantification. Frozen samples were shipped to Michigan State University Mass Spectrometry and Metabolomics Core (MSU MSMC, East Lansing, MI; protocol ID: MSU_MSMC_010a) for analysis by gas chromatography/mass spectrometry (GC/MS). SCFA concentrations were calculated by normalization to standards, then normalized by baseline levels detected in each media (MiPro, low-CF-MiPro and median-CF-MiPro). Finally, these concentrations were normalized to the CFU/mL calculated in that exact sample to account for differences in microbial density. SCFA concentrations over time were displayed using ggplot2 (version 3.4.2) and linear mixed models were used for statistical regression analysis to account for repeated measures within sample source (lme4 version 1.1.34). For each sample, beta diversity was calculated by Bray-Curtis distance, and multidimensional scaling ordination was performed via PCA plots. Significant differences in beta diversity were tested by permutational analysis of variance (PERMANOVA) (vegan version 2.6.4).

## Acknowledgement

We thank Dartmouth Translational Research Core for facilitating retrieval and storage of stool and colonoscopy samples, and the individuals who generously donated samples. We thank Melissa Carmichael for her technical assistance. We also thank Ali Kohan and Mark Sundrud for helpful comments and feedback. This work was supported by CFF grant OTOOLE22G0 and NIHES033988 to GAO, CFF grant MADAN18A0 to JCM, NIH/T32HL134598 to KEB, National Science Foundation award DGE-2125733 and the Innovation PhD Program to RAV, with additional support from DartCF (NIH grant P30-DK117469) and the Cystic Fibrosis Foundation Research Development Program (STANTO19R0).

