## Supplemental Figures for "An In Vitro Medium for Modeling Gut Dysbiosis Associated with Cystic Fibrosis"

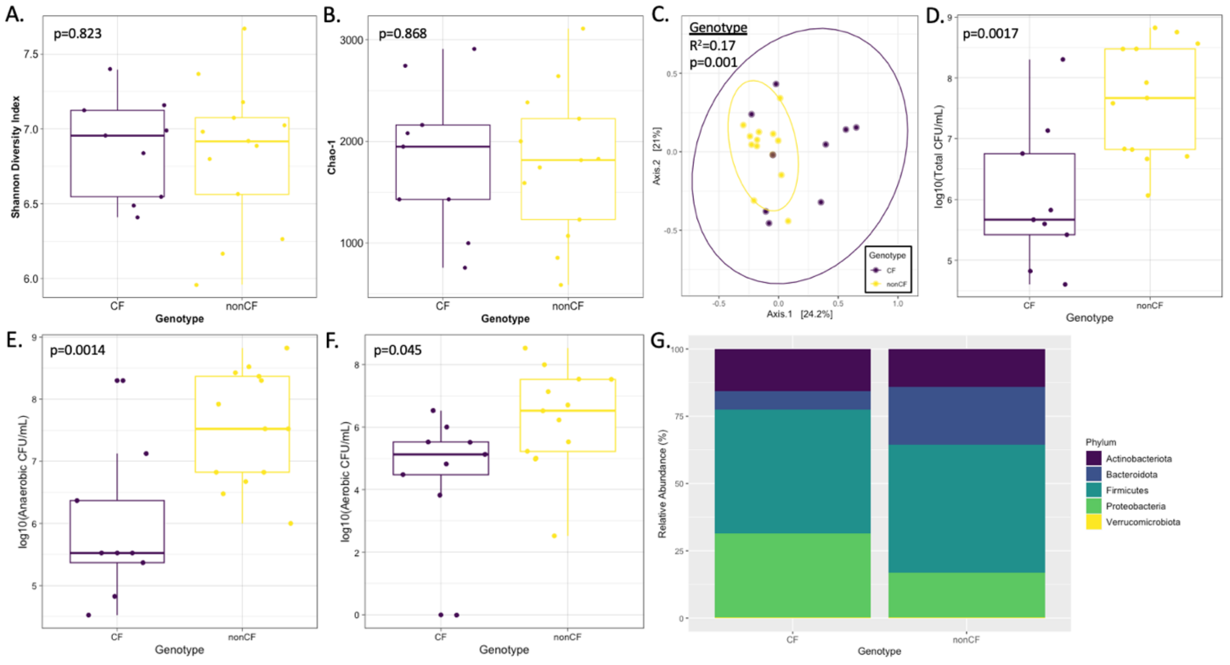


**Supplemental Figure 1. Description of CF and nonCF stool or colonoscopy aspirates used as a source of the inoculum in the studies presented here.** Alpha diversity of CF and nonCF stool and/or colonoscopy aspirates using (**A**) Shannon Diversity Index (SDI) or (**B**) Chao-1 distance. A linear model from the R package stats was used to test whether SDI or Chao-1 changed significantly with genotype. Neither SDI nor Chao-1 are significantly affected by genotype. (**C**) Bray-Curtis beta diversity was calculated for each sample and displayed on a principal coordinate analysis (PCA) plot, colored by genotype. The first two components account for 45.2% of total variance. Significant differences in beta diversity due to genotype were tested by PERMANOVA (p=0.001). (**D**) Sum CFU/mL were calculated following growth on blood sheep agar at (**E**) 0% or (**F**) 21% oxygen. A linear model from the R package stats was used to test whether CFU/mL from each oxygen tension changed significantly with genotype. CF samples culture less CFU/mL at 0% (**E**) and 21% oxygen (**F**), thus less total CFU/mL compared to nonCF samples (**D**). **(G)** Relative abundance of the top five phyla, legend to the right, of CF and nonCF samples. In CF samples, Proteobacteria (green) is increased and Bacteroidota (blue) is decreased.

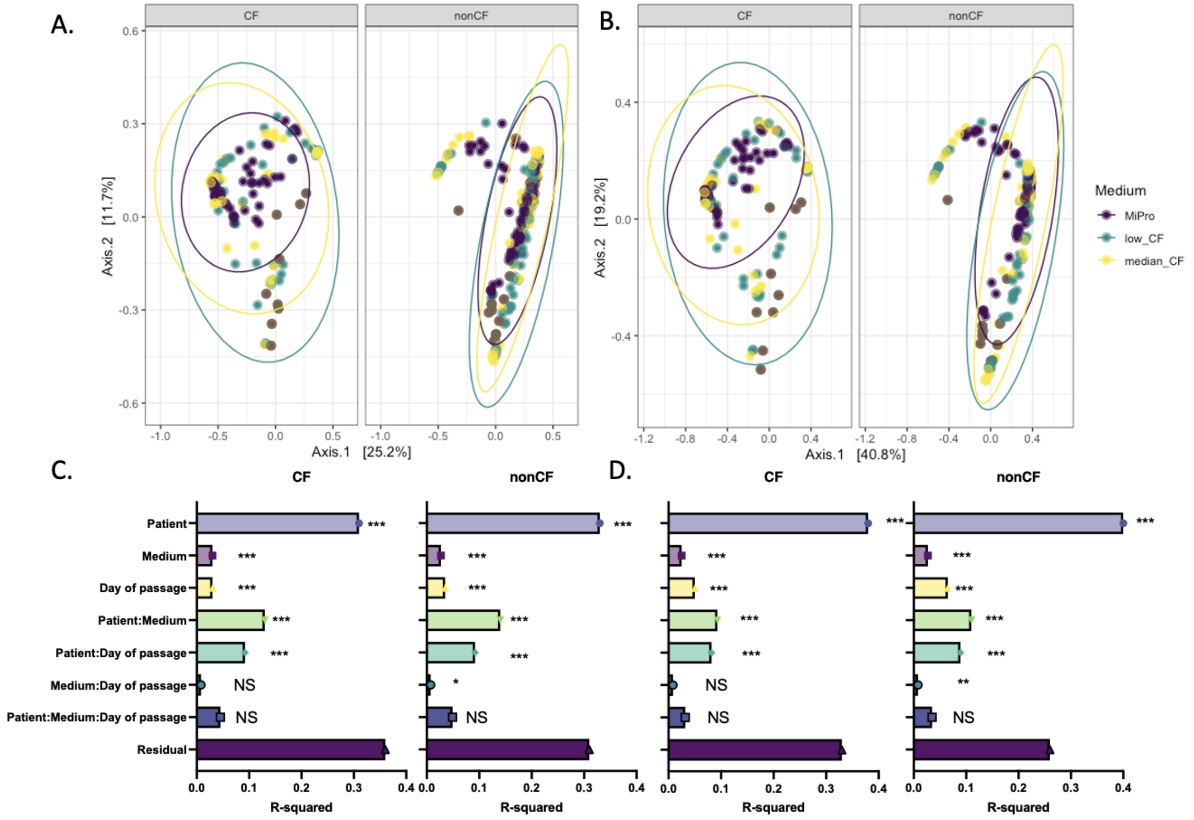


**Supplemental Figure 2. Jaccard and Morisita-Horn beta diversity of CF and nonCF samples cultured through MiPro and CF-MiPro.** (**A**) Jaccard and (**B**) Morisita-Horn beta diversity was calculated for each sample and displayed on PCA plots, faceted by genotype and colored by medium. The first two components account for (**A**) 36.9% and (**B**) 60% of total variance. Statistical differences in (**C**) Jaccard and (**D**) Morisita-Horn beta diversity were tested by PERMANOVA, with metadata included in the model. R-squared values for all data included, as well as potential interactions, are plotted with significance codes indicated (NS: non-significant, *: p<0.05, **: p<0.01, ***: p<0.001). “Residual” indicates variation unexplained by the model.

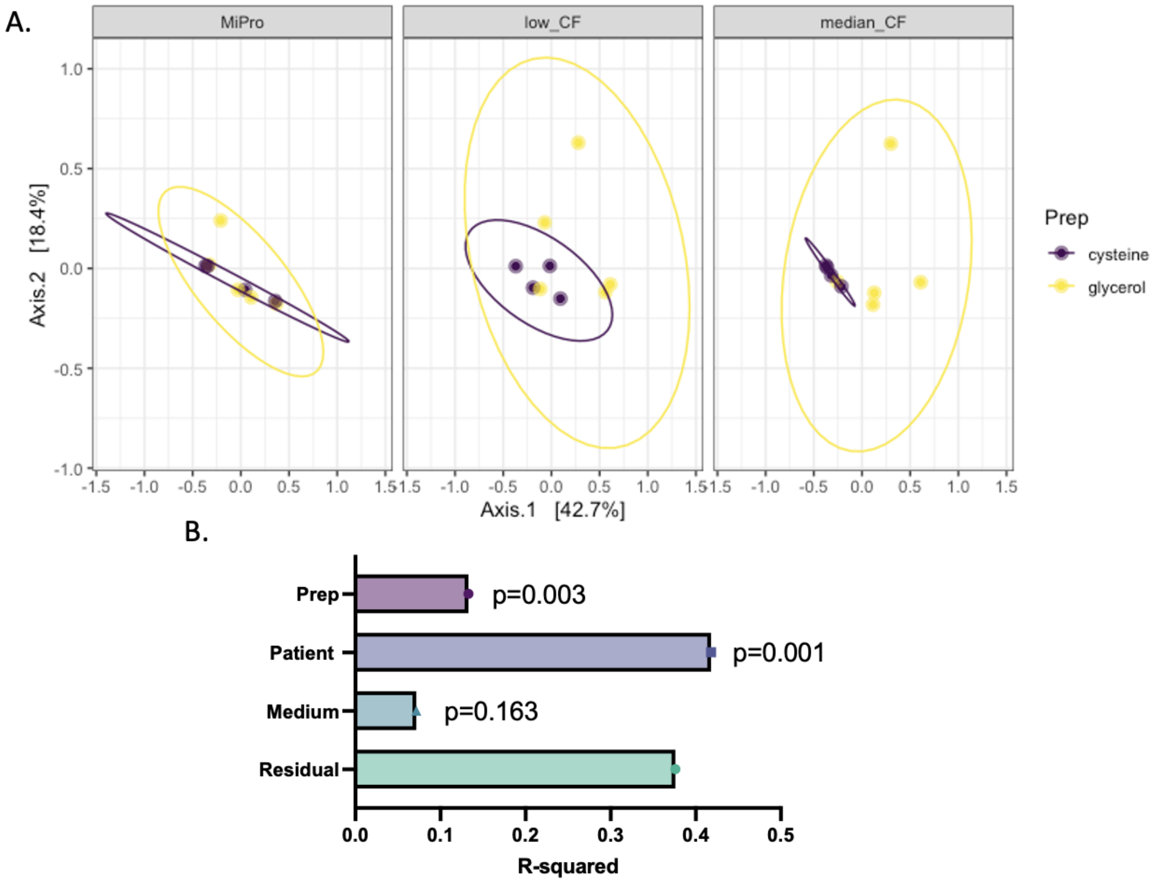


**Supplemental Figure 3. Bray-Curtis beta diversity of CF Day 5 samples homogenized in glycerol or L-cysteine.** (**A**) Bray-Curtis beta diversity was calculated for each sample and displayed on PCA plots, faceted by medium and colored by "Prep". "Prep" signifies the solution in which raw stool was homogenized on Day 0: PBS supplemented with 7.15% glycerol or 10 mM L-cysteine. The first two components account for 61.1% of total variance. Statistical differences in Bray-Curtis beta diversity were tested by PERMANOVA, with metadata included in the model. R-squared values for all data included are plotted with p-values indicated. "Residual" indicates variation unexplained by the model.

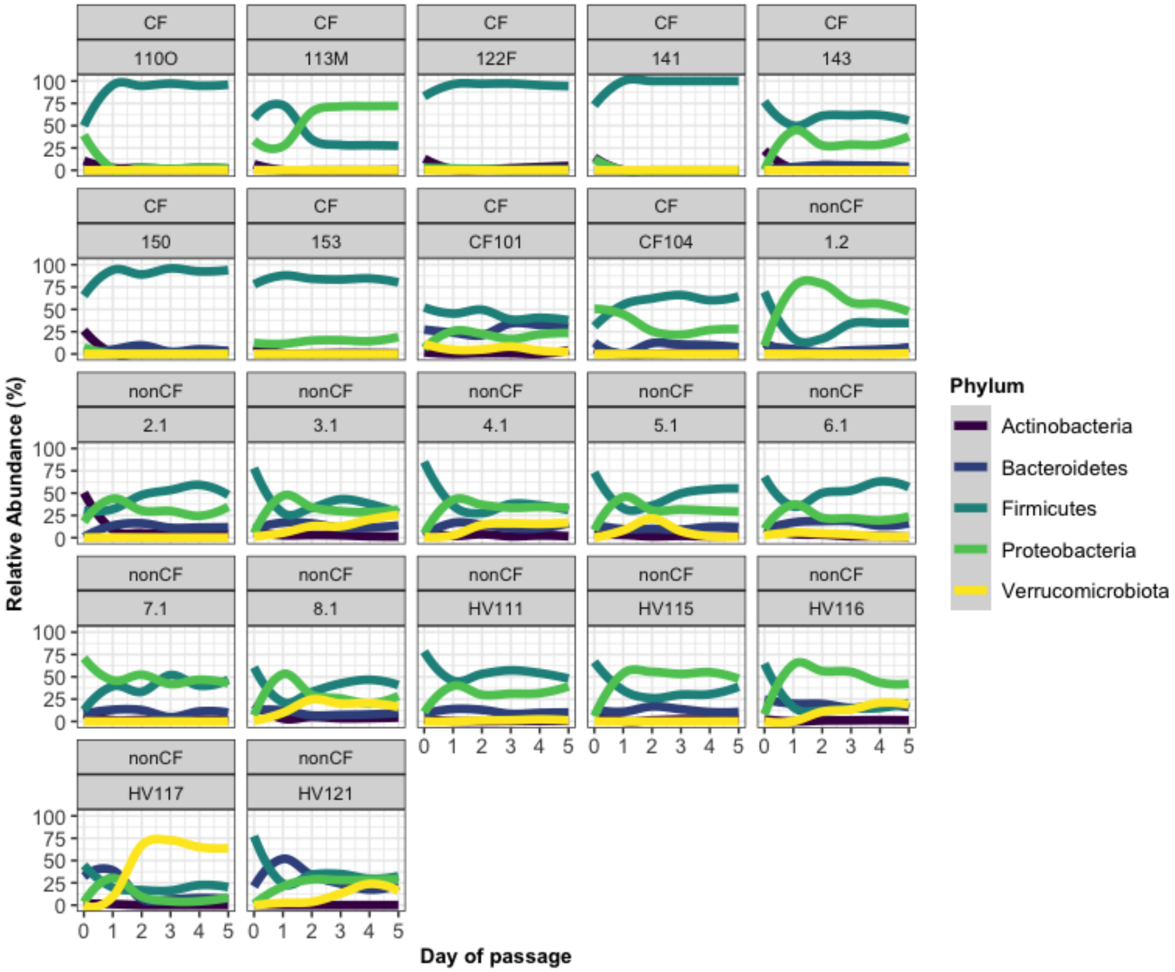


**Supplemental Figure 4. Intra-person variation of microbial relative abundance changes in MiPro.** Day of passage is graphed versus relative abundance of the indicated taxa for each sample, and a linear plot was used to visualize overall changes in microbial relative abundance at the phylum level during in vitro passages in MiPro. The legend indicates the taxonomic assignment for each panel. Taxonomical shifts occur largely at Day 1 and are not restricted to a specific genotype, sample type (stool or colonoscopy aspirate), homogenization protocol (“Prep”: glycerol or cysteine) or sequencing batch.

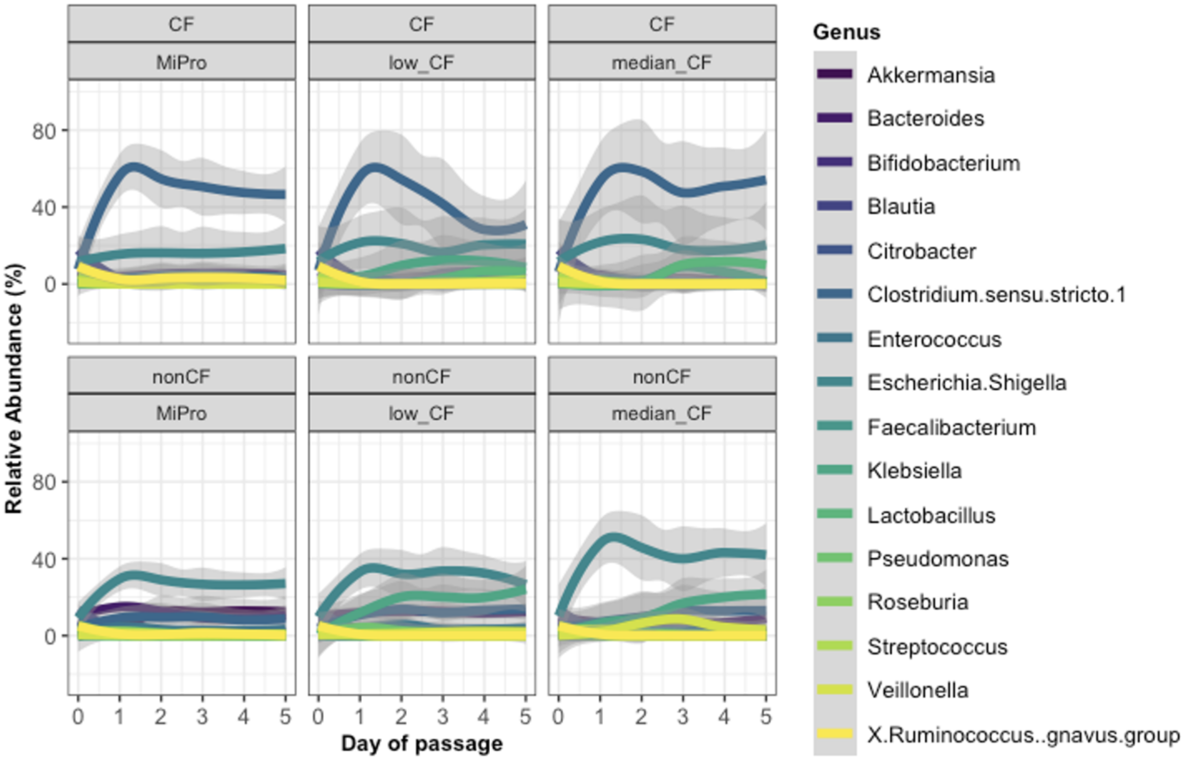


**Supplemental Figure 5. Change in genera over time.** Day of passage is graphed versus relative abundance of the indicated taxa for each sample, and a linear plot was used to visualize overall changes in microbial relative abundance at the genus level across all media conditions. Samples originating from a CF donor are displayed in the top panels, and samples originating from a nonCF donor are displayed in the bottom panels. The legend indicates the taxonomic assignment for each panel.
