## Supplemental Tables for "An In Vitro Medium for Modeling Gut Dysbiosis Associated with Cystic Fibrosis"

**Table S3. Relative abundance of major phyla across media conditions.**

| *Phylum* | *CF* | | | | *nonCF* | | | |
|---|---|---|---|---|---|---|---|---|
| | **Raw*** | **MiPro**** | **Low-CF-MiPro** | **Median-CF-MiPro** | **Raw** | **MiPro** | **Low-CF-MiPro** | **Median-CF-MiPro** |
| **Actinobacteria** | 12.3 | 1.47 | 1.85 | 1.73 | 10.5 | 2.03 | 2.41 | 2.93 |
| **Bacteroidota** | 4.89 | 5.19 | 2.93 | 1.86 | 12.7 | 13.5 | 13.5 | 8.32 |
| **Firmicutes** | 63 | 74.4 | 60.1 | 72.2 | 62.3 | 36.1 | 22.2 | 23.7 |
| **Proteobacteria** | 18.3 | 18.3 | 35.1 | 24.2 | 12.3 | 36.9 | 60.8 | 64.7 |
| **Verrucomicrobiota** | 1.27 | 0.543 | 0.0088 | 0.0113 | 0.441 | 10.8 | 0.727 | 0.0828 |

*"Raw" represents the average relative abundance of the indicated phylum in uncultured colonoscopy aspirate or homogenized stool.
**Each medium condition represents the average relative abundance of the indicated phylum across all days of passage (1-5) in that medium.

**Table S4. Mixed linear model of SDI from CF samples cultured in MiPro over five days.**

| | Shannon | | |
|---|---|---|---|
| *Predictors* | *Estimates* | *CI* | *p* |
| (Intercept) | 6.58 | 6.28 – 6.88 | **<0.001** |
| Day of passage | -0.03 | -0.08 – 0.03 | 0.293 |
| **Random Effects** | | | |
| $\sigma^2$ | 0.11 | | |
| $\tau_{00\ Patient}$ | 0.14 | | |
| ICC | 0.55 | | |
| $N_{Patient}$ | 9 | | |
| Observations | 54 | | |
| Marginal $R^2$ / Conditional $R^2$ | 0.009 / 0.558 | | |

**Table S5. Mixed linear model of SDI from nonCF samples cultured in MiPro over five days.**

| | Shannon | | |
|---|---|---|---|
| *Predictors* | *Estimates* | *CI* | *p* |
| (Intercept) | 6.80 | 6.58 – 7.03 | **<0.001** |
| Day of passage | 0.01 | -0.02 – 0.04 | 0.411 |
| **Random Effects** | | | |
| $\sigma^2$ | 0.05 | | |
| $\tau_{00\ \text{Patient}}$ | 0.14 | | |
| ICC | 0.73 | | |
| $N_{\text{Patient}}$ | 13 | | |
| Observations | 78 | | |
| Marginal $R^2$ / Conditional $R^2$ | 0.002 / 0.731 | | |

**Table S6. Mixed linear model of Chao1 from CF samples cultured in MiPro over five days.**

| | Chao 1 | | |
|---|---|---|---|
| *Predictors* | *Estimates* | *CI* | *p* |
| (Intercept) | 1464.40 | 1041.55 – 1887.24 | **<0.001** |
| Day of passage | -37.31 | -101.63 – 27.00 | 0.249 |
| **Random Effects** | | | |
| $\sigma^2$ | 161480.77 | | |
| $\tau_{00\ Patient}$ | 314292.17 | | |
| ICC | 0.66 | | |
| $N_{Patient}$ | 9 | | |
| Observations | 54 | | |
| Marginal $R^2$ / Conditional $R^2$ | 0.009 / 0.664 | | |

**Table S7. Mixed linear model of Chao1 from nonCF samples culture in MiPro over five days.**

| | Chao 1 | | |
|---|---|---|---|
| *Predictors* | *Estimates* | *CI* | *p* |
| (Intercept) | 1761.01 | 1416.53 – 2105.49 | **<0.001** |
| Day of passage | 2.85 | -33.17 – 38.87 | 0.875 |
| **Random Effects** | | | |
| $\sigma^2$ | 74339.07 | | |
| $\tau_{00\ Patient}$ | 349621.92 | | |
| ICC | 0.82 | | |
| $N_{Patient}$ | 13 | | |
| Observations | 78 | | |
| Marginal $R^2$ / Conditional $R^2$ | 0.000 / 0.825 | | |

**Table S8. Relative abundance of top 10 families across media conditions.**

| *Family* | *CF* | | | | *nonCF* | | | |
|---|---|---|---|---|---|---|---|---|
| | **Raw*** | **MiPro**** | **Low-CF-MiPro** | **Median-CF-MiPro** | **Raw** | **MiPro** | **Low-CF-MiPro** | **Median-CF-MiPro** |
| **Akkermansiaceae** | 1.29 | 0.543 | 0.00864 | 0.0113 | 0.399 | 10.7 | 0.716 | 0.0674 |
| **Bacteroidaceae** | 3.41 | 4.94 | 2.86 | 1.42 | 9.9 | 11.6 | 12.2 | 7.06 |
| **Bifidobacteriaceae** | 6.87 | 0.268 | 1.44 | 1.37 | 7.9 | 0.723 | 1.61 | 2.65 |
| **Clostridiaceae** | 4.61 | 54.6 | 43.5 | 54.6 | 3.82 | 9.6 | 12.8 | 11.1 |
| **Enterobacteriaceae** | 14.5 | 17.6 | 34.8 | 24.1 | 8.93 | 32.2 | 54.9 | 62.5 |
| **Lachnospiraceae** | 42.9 | 8.58 | 1.6 | 0.398 | 39.2 | 8.1 | 0.948 | 0.429 |
| **Lactobacillaceae** | 1.4 | 0.233 | 10.9 | 14.7 | 0.323 | 0.0923 | 0.311 | 2.73 |
| **Pseudomonadaceae** | 3.05 | 0.0432 | 0.0438 | 0.0201 | 0.317 | 0.492 | 2.18 | 0.258 |
| **Ruminococcaceae** | 1.57 | 0.236 | 0.0196 | 0.00318 | 7.77 | 0.277 | 0.0623 | 0.0212 |
| **Veillonellaceae** | 1.91 | 0.312 | 0.074 | 0.0142 | 0.738 | 1.41 | 2.07 | 4.38 |

*"Raw" represents the average relative abundance of the indicated phylum in uncultured colonoscopy aspirate or homogenized stool.
**Each medium condition represents the average relative abundance of the indicated phylum across all days of passage (1-5) in that medium.

**Table S9. Relative abundance of top 15 genera across media conditions.**

| *Genus* | *CF* | | | | *nonCF* | | | |
|---|---|---|---|---|---|---|---|---|
| | Raw* | MiPro** | Low-CF-MiPro | Median-CF-MiPro | Raw | MiPro | Low-CF-MiPro | Median-CF-MiPro |
| ***Akkermansia*** | 1.27 | 0.543 | 0.00864 | 0.0113 | 0.399 | 10.7 | 0.716 | 0.0673 |
| ***Bacteroides*** | 3.41 | 4.94 | 2.86 | 1.42 | 9.9 | 11.6 | 12.2 | 7.06 |
| ***Bifidobacterium*** | 6.86 | 0.267 | 1.43 | 1.37 | 7.9 | 0.706 | 1.6 | 2.65 |
| ***Blautia*** | 18.4 | 1 | 0.162 | 0.31 | 13.9 | 1.47 | 0.233 | 0.199 |
| ***Citrobacter*** | 0 | 0.2 | 3.5 | 0.00353 | 0.011 | 0.534 | 0.579 | 0.0121 |
| ***Clostridium sensu stricto 1*** | 4.58 | 53.3 | 43.5 | 54.6 | 3.76 | 9.2 | 12.8 | 11.1 |
| ***Enterococcus*** | 0.181 | 3.83 | 2.36 | 1.48 | 0.58 | 3.58 | 3.91 | 2.88 |
| ***Escherichia/Shigella*** | 11.4 | 16.6 | 20.7 | 20.3 | 7.99 | 28.7 | 32.9 | 45.9 |
| ***Faecalibacterium*** | 0.611 | 0.189 | 0.00125 | 0.0016 | 5.95 | 0.00716 | 0.0146 | 0.0101 |
| ***Klebsiella*** | 2.89 | 0.405 | 8.93 | 3.36 | 0.711 | 1.41 | 18.9 | 15.1 |
| ***Lactobacillus*** | 0.456 | 0.0718 | 3.36 | 6.33 | 0.143 | 0.0205 | 0.0206 | 0.951 |
| ***Pseudomonas*** | 3.05 | 0.0429 | 0.0438 | 0.0201 | 0.317 | 0.492 | 2.17 | 0.258 |
| ***Ruminococcus gnavus group*** | 9.57 | 2.51 | 0.167 | 0.00857 | 5.28 | 0.627 | 0.0464 | 0.0605 |
| ***Streptococcus*** | 5.25 | 1.33 | 0.924 | 0.404 | 2 | 1.21 | 0.321 | 1.28 |
| ***Veillonella*** | 1.07 | 0.0147 | 0.0024 | 0.0142 | 0.562 | 0.461 | 1.65 | 4.36 |

*"Raw" represents the average relative abundance of the indicated phylum in uncultured colonoscopy aspirate or homogenized stool.

**Each medium condition represents the average relative abundance of the indicated phylum across all days of passage (1-5) in that medium.
